# Frequency Comb Optoacoustic Tomography

**DOI:** 10.1101/2021.05.12.443808

**Authors:** Antonios Stylogiannis, Ludwig Prade, Sarah Glasl, Qutaiba Mustafa, Christian Zakian, Vasilis Ntziachristos

## Abstract

Optoacoustics (OA) is overwhelmingly implemented in the Time Domain (TD) to achieve a high Signal-to-Noise-Ratio (SNR). Implementations in the Frequency Domain (FD) have been proposed, but have not offered competitive advantages over TD methods to reach high dissemination. It is therefore commonly believed that the TD represents the optimal way of performing optoacoustics. Here, we introduce a novel optoacoustic concept based on frequency comb and theoretically demonstrate its superiority to the TD. Then, using recent advances in laser diode illumination, we launch Frequency Comb Optoacoustic Tomography (FCOT), at multiple wavelengths, and experimentally demonstrate its advantages over TD methods in phantoms and *in-vivo*. We demonstrate that FCOT optimizes the SNR of spectral measurements over TD methods by benefiting from signal acquisition in the TD and processing in the FD, and that it reaches the fastest multi-spectral operation ever demonstrated in optoacoustics while reducing performance compromises present in TD systems.

## Introduction

Generation of optoacoustic (OA) signals requires illumination with energy transients^1,2^. Time Domain (TD) implementations offer large energy transients by means of nanosecond duration light pulses^3–6^, optimizing the signal-to-noise ratio (SNR) and making TD the domain of choice in optoacoustics^7–9^. TD optoacoustic imaging records the time-of-flight of ultrasound waves (US) at multiple locations on the surface of the interrogated object and, using mathematical inversion, converts these measurements to maps of optical absorption^10^.

Other imaging modalities such as optical coherence tomography (OCT) or magnetic resonance imaging (MRI) were originally demonstrated in the TD, but have tremendously benefited from switching operation to the Frequency Domain (FD)^11,12^. The advantages of FD were so fundamental that all OCT or MRI implementations offered today operate in the FD. However, FD-domain optoacoustic implementations using intensity-modulated light^13–15^ provide up to six orders of magnitude^16^ smaller energy transients compared to ultrashort pulses, drastically reducing the SNR^17–19^. Moreover, we have recently shown^20^ the need to capture many discrete frequencies for image generation, which may lead to complex detection (demodulation) schemes ^20,21^. Therefore, despite the higher duty cycles achieved over the TD ^16,20–23^, FD optoacoustics has offered little impact in the imaging field.

Frequency chirp has also been investigated as a hybrid TD-FD method, by modulating light at a continuously varying frequency^22,24^, thus encoding time in frequency. Detection is carried out in the TD using time-correlation techniques. Similar to FD methods however, the use of sine waves limits the achieved SNR, restricting the use of chirp approaches to experimental investigations.

We propose Frequency Comb (FC) operation to categorically improve upon TD performance, while minimizing FD disadvantages. FC is effectively the reverse implementation of chirp optoacoustics, using a train of discrete pulses, similar to the ones employed in TD optoacoustics, but processing the resulting signals in the FD, representative of an FD system at multiple discrete frequencies^20^. As shown herein, FC illumination allows concurrent illumination at multiple wavelengths without increasing the imaging time and yielding an SNR gain that increases by the square root of the number of wavelengths N employed, over TD systems.

Following theoretical considerations, we hypothesized that overdriving laser diodes with FC pulse trains could lead to high quality optoacoustic imaging that could demonstrate benefits over TD implementations. We have shown that overdriven laser diodes yield inexpensive optoacoustic illumination at > 130nJ per pulse at repetition rates at the hundreds of KHz^25^. Based on this illumination ability, we introduce a **Frequency Comb Optoacoustic Tomography (FCOT)** system (**Suppl. Fig.1**), implemented with 4 concurrently pulsing wavelengths at 6.8ns pulse width, using custom made laser diode drivers and Arbitrary Waveform Generators (AWG). Each wavelength is pulsed at a slightly different repetition rate (**see methods**). Detection was performed with a 50MHz central frequency and 110% bandwidth ultrasound transducer (UST), imparting mesoscopic operation^26^. We show concurrent multi-wavelength imaging of lymphatic and microvascular dynamics in mice at high SNRs, offering the fastest multi-wavelength illumination ever achieved in the field of optoacoustics and confirming spectral performance that improves upon TD implementations.

## Results

### Frequency Comb Optoacoustics using a single excitation wavelength

In conventional TD operation, a square light pulse of duration t_p_ (Fig.1a) yields a continuous frequency spectrum in the FD, via the Fourier Transform, with the first node at the 1/t_p_ frequency. In FD operation, light modulated by a sine wave in the TD (Fig.1b) yields a single discrete frequency in the FD. FC modulation instead considers a train of pulses (Fig.1c) with a pulse width of t_p_ and a repetition rate f_rep_. The Fourier Transform of this pulse train yields many discrete frequencies with an envelope (Fig. 1c, right) identical to the continuous spectrum of a single pulse with duration t_p_ (Fig.1a, right). The discrete frequencies of the pulse train are harmonics of the fundamental repetition rate, f_rep_, i.e. integer multiples of f_rep_. The frequency spacing and the envelope observed in the FD is determined by the repetition rate of the pulse train (Fig.1d) and the pulse width (Fig.1e) respectively.

**Figure 1.**
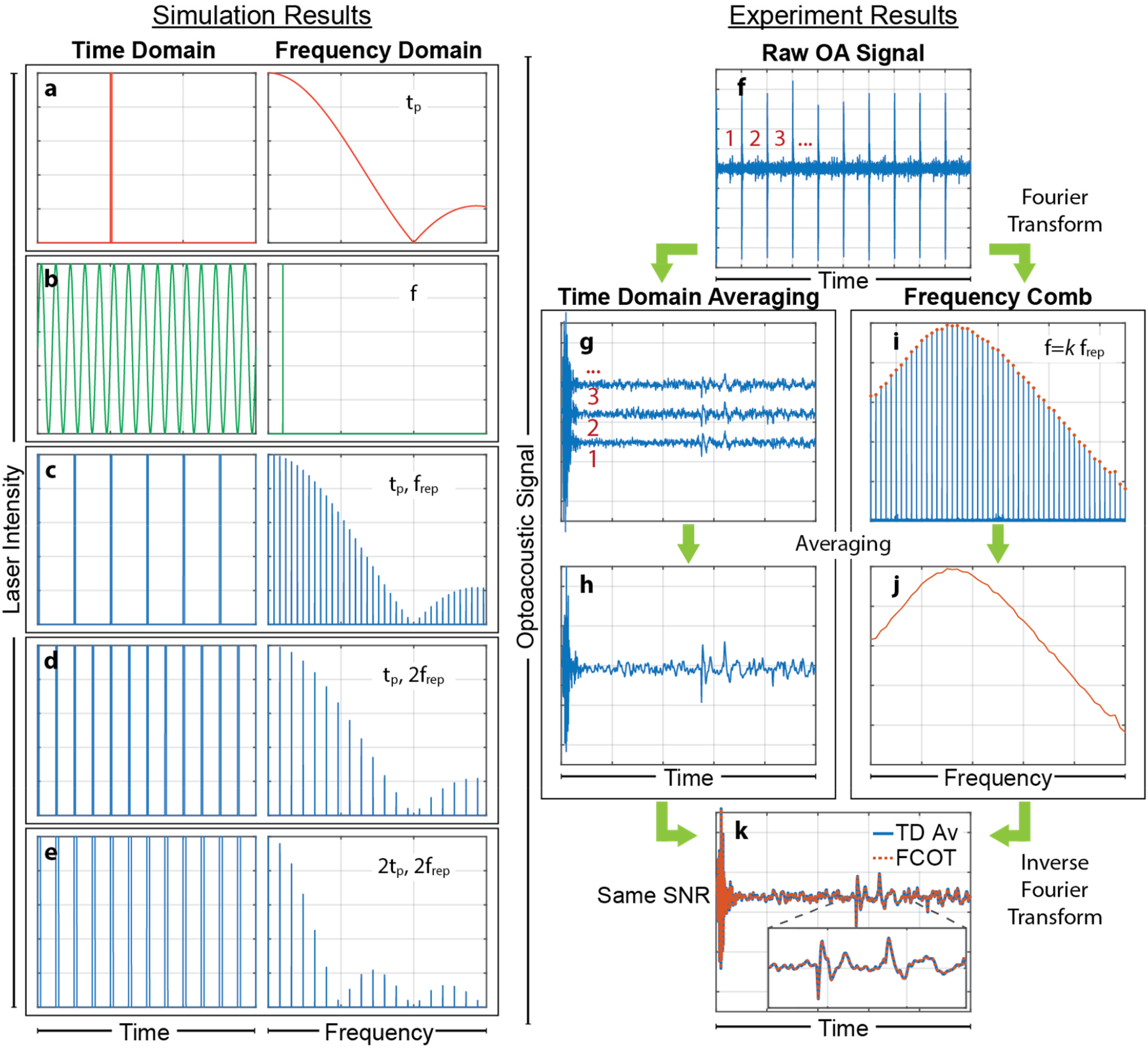
The signal processing algorithms used in Frequency Comb Optoacoustics at a single wavelength. **a)** A single excitation light pulse of t_p_ duration in TD (left) and a continuous spectrum of frequencies in FD (right). **b)** A sine wave of frequency f in TD and FD, continuous wave in TD and a single discrete peak in FD. **c)** A train of pulses with pulse duration t_p_ and repetition rate f_rep_ in TD and FD. Many discrete pulses in TD and many discrete frequencies in FD with the same envelope as a single pulse of t_p_ duration in **a). d)** A train of pulses with pulse duration t_p_ and repetition rate 2f_rep_ in TD and FD. Twice as many pulses in TD but half the number of discrete frequencies in FD compared to **c). e)** A train of pulses with pulse duration 2t_p_ and repetition rate 2f_rep_ in TD and FD. As many pulses in TD and discrete frequencies in FD as in **d)** but now following a different envelope than **a)** or **c). f)** The raw optoacoustic signal recorded using a pulse train like **c)** for example. **g**,**h)** present the normal averaging in TD. **g)** The train of pulses is split in sections of period T=1/f_rep_ which are averaged **(h)** point by point. **i**,**j)** The Frequency Comb processing of the same signal. **i)** The Fourier transform of the raw optoacoustic signal **(f)** with many discrete frequencies that are all harmonics (k*f_rep_ with k positive integer) of the base repetition rate f_rep_. In FD we choose only the harmonics of f_rep_ and discard all the other frequencies that contain only noise **(j). k)** By performing the inverse Fourier Transform in **j)** we recover the TD signal that matches perfectly with the one in **h)**.

Experimental validation of the FC scheme was performed by exciting a black varnish layer on a petri dish at 445nm, using a pulse train of t_p_=6.8ns and f_rep_=200kHz delivering 189nJ per pulse (**Fig.1f,g**). Averaging in TD (Fig.1g-h) increases the SNR by a factor of sqrt(N_p_), where N_p_ is the number of pulses in the pulse train. In contrast, the proposed FC method selects the fundamental frequency f_rep_ and its harmonics, *k*\*f_rep_ (Fig.1i) by performing the following operation (see also Suppl. materials):

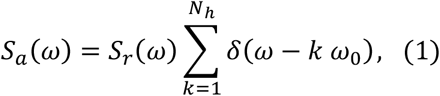

where *S*_*a*_(*ω*) is the Fourier Transform of the averaged signal, *S*_*r*_(*ω*) the Fourier Transform of the recorded signal (Fig.1f), *N*_*h*_ the number of the harmonics in the recorded bandwidth and *ω*_0_ = 2 π/*T*. This operation selects only the harmonics of f_rep_ and filters out frequencies that do not contain signal to increase the SNR by the same factor sqrt(N_p_) as in the TD (Fig.1j). The signal in Fig.1j is the Fourier Transform of the signal in Fig.1h, with the two signals matching perfectly (Fig.1k). This analysis confirms that FC illumination results in a practical generation of multiple discrete frequencies, required for accurate FD operation, offering an SNR that is equivalent to the TD when a single wavelength is used. Next, we show however that FC offers advantages over TD when multiple wavelengths are employed.

### Frequency Comb Optoacoustics using multiple excitation wavelengths

To demonstrate the FC advantage over the TD (Fig.2) we plotted the single-wavelength excitation pattern in the TD and its power spectrum in Fourier space (Fig.2a) to serve as reference for the analysis that follows. The pulse train shown has a period T that corresponds to a repetition rate f_rep_=1/T, a total number of pulses N_p_ and an acquisition time t_acq_=N_p_*T. The period T defines the maximum depth of view DoV=v_s_*T that can be achieved for the pulse train selected, where v_s_ is the speed of sound. TD wavelength multiplexing is performed using wavelength interleaving, or time sharing. However, when increasing the number of wavelengths in the TD, at least one of the following three parameters has to be compromised: the DoV, the number of pulses in the pulse train and therefore the SNR for each wavelength, or the total acquisition time. Fig.2b shows how the DoV is reduced when using four wavelengths at a given total acquisition time. The different wavelengths excite the tissue using the same repetition rate f_rep_ but with a time shift t_sh_ between each wavelength (Fig.2a), given by t_sh_=T/N, where N is the number of wavelengths. The result is a reduction of the time between subsequent pulses, limiting the DoV available to each wavelength by a factor N. Alternatively, it is possible to retain the original DoV by dropping the repetition rate for each wavelength, j, to f_rep,j_=f_rep_/N and the number of pulses per wavelength to N_p_/N (Fig.2c), resulting however in an SNR reduction by a factor of sqrt(N). A third alternative, retaining the original DoV and SNR, is to prolong the acquisition time by a factor N (Fig.2d).

**Figure 2.**
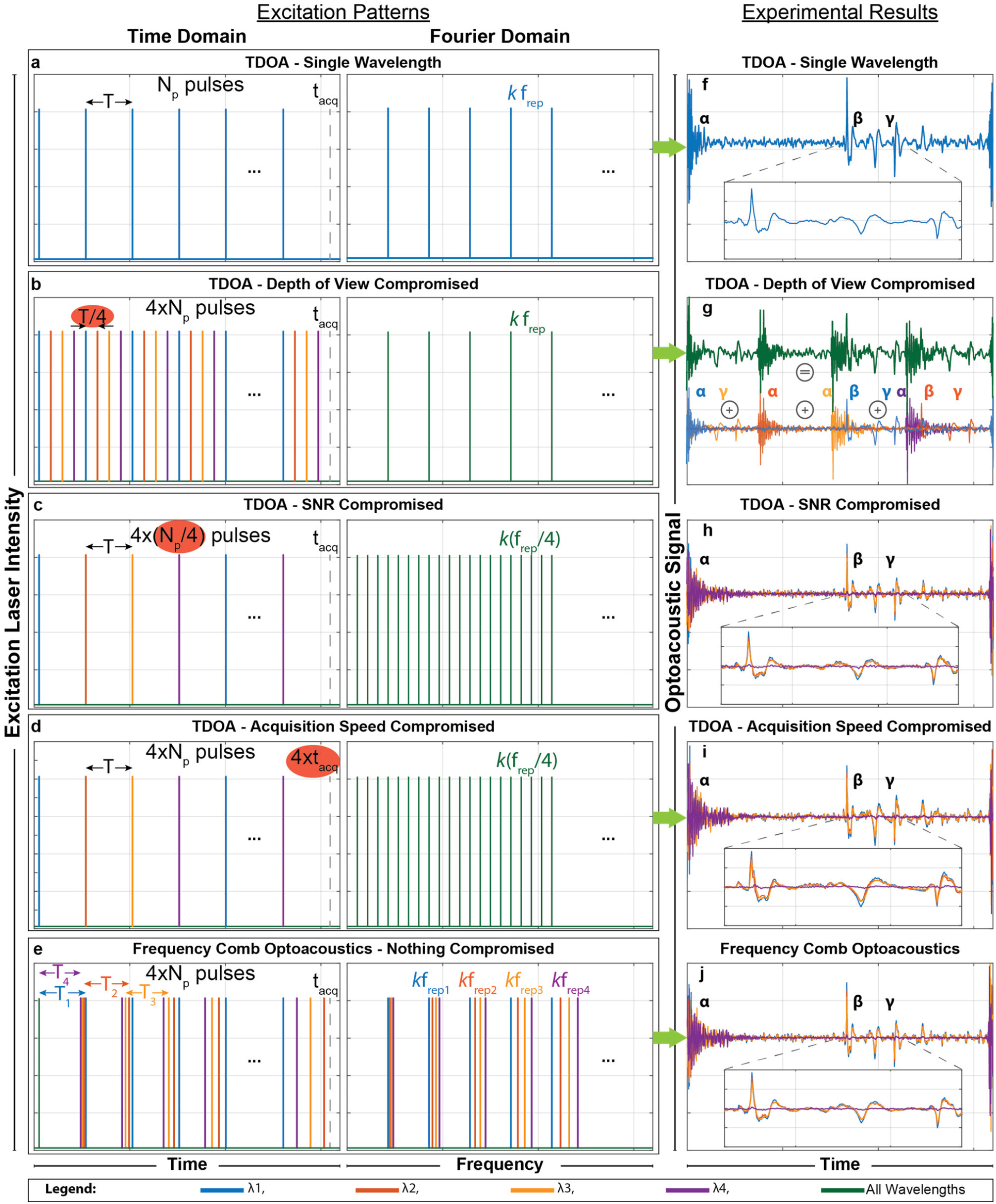
FCOT advantages at multiple wavelengths. **a)** A pulse train of a single wavelength with period T and repetition rate f_rep_=1/T, with N_p_ pulses and an acquisition time t_acq_ as it appears in the time domain (left) and the Fourier domain (right). **(b)-(d)** Multiple wavelengths excitation in TD Optoacoustic. **(b)** The excitation pattern of 4 wavelengths emitting at the same repetition rate f_rep_ with a time shift equal to T/4, N_p_ pulses for all 4 wavelengths and t_acq_ acquisition time. In this case the depth of view of each wavelength is compromised. **(c)** The excitation pattern of 4 wavelengths with f_rep_/4 repetition rate, N_p_/4 pulses for each wavelength and t_acq_ acquisition time. In this case the SNR is compromised. **(d)** The excitation pattern of 4 wavelengths with f_rep_/4 repetition rate, N_p_ pulses per wavelength but with 4t_acq_ acquisition time. In this case the acquisition time is compromised. **(e)** Frequency Comb Optoacoustic excitation where all 4 wavelengths have different repetition rates f_rep1_, f_rep2_, f_rep3_, f_rep4_ between them, N_p_ pulses for each wavelength and t_acq_ acquisition time. **(f)-(j)**The Optoacoustic signal recorded by a layer of black varnish on a petri dish from the excitation patterns in **(a)**-**(e)** respectively. **(f)** The optoacoustic signal from a single wavelength. **α**, the electromagnetic interference from the laser diode circuitry when triggered, **β** the optoacoustic signal from the black varnish, **γ** the reflection of the optoacoustic signal in the petri dish or in the acoustic lens of the UST. **(g)** The optoacoustic signal from the excitation pattern (b) for all wavelengths (green line) that is the sum of the optoacoustic signal from each wavelength separately (bottom line). The laser interference, optoacoustic signal and its reflections (**α, β, γ**) for each wavelength are indicated. The depth of view for each wavelength is drastically reduced. **(h)** The optoacoustic signal from the excitation pattern **(c)**. The SNR of the optoacoustic signal of each wavelength is reduced. **(i)** The optoacoustic signal from the excitation pattern **(d)**. The SNR and depth of view of the optoacoustic signal of each wavelength are maintained but the acquisition time is increased. **(j)** The optoacoustic signal where Frequency Comb Optoacoustic is used. Signal from all 4 wavelengths has been recovered without any cross-talk between the lasers and correctly co-registered in time without compromising depth of view, SNR or acquisition time.

In contrast to TD, FCOT employs a small frequency shift δf for the repetition rate of each wavelength (Fig.2e; time), whereby δf<<f_rep_. Each wavelength has a different repetition rate which results in a slightly different effective DoV; however, since δf<<f_rep_, this difference is insignificant. The power spectrum of the FCOT excitation pattern (Fig.2e; right) shows the appearance of harmonics from the fundamental repetition rate for each wavelength. The repetition rate of the first laser, f_rep,1_, can be selected as the reference repetition rate, with the repetition rate of the remaining lasers given by f_rep,j_=f_rep,1_+(j-1)*δf. FCOT thus recovers the OA signals by evaluating equation (1) with the corresponding ω_0,j_=2πf_rep,j_ for each wavelength. Consequently, using frequency separation, FCOT can multiplex different wavelengths without compromising the DoV, SNR or acquisition time. With the signal processed in the FD, the frequency resolution is defined by df=1/t_acq_, where t_acq_=N_p_/f_rep1_, meaning that frequencies that differ by less than df cannot be resolved. In order to recover the OA signal from all wavelengths, all harmonics from all lasers that lie in the UST detection bandwidth should therefore be spaced at a distance exceeding the frequency resolution df, imposing a minimum number of pulses N_p,min_ that depends on the UST bandwidth limits (f_low_ and f_high_), N and f_rep,1_ (see **suppl. material**). The parameters N, f_low_, f_high_, f_rep,1_ and the chosen number of pulses, N_p_, define the range of small frequency shifts (δf), between δf_min_ and δf_max,_, required to recover the signal of each wavelength without losses. A δf value larger than δf_min_ ensures that harmonics of laser j and j+1, which lie on the lower end of the UST bandwidth, are well-resolved, while a δf value smaller than δf_max_ ensures that harmonics of laser 1 and N, which lie on the upper end of the UST bandwidth, are well-resolved.

### FCOT outperforms TDOA in SNR, DoV or imaging speed

To experimentally demonstrate the advantages of FC operation in multi-wavelength excitation compared to TD optoacoustics, we employed FCOT with four wavelengths at 445, 465, 638 and 808 nm, named wavelengths 1, 2, 3 and 4, respectively. The resulting optoacoustic signal at wavelength 1, f_rep,1_=200kHz and N_p_=200 (Fig.2f) attained t_acq_=1ms and DoV=7.5mm. Electromagnetic interference from the laser diode (LD) driving circuit is indicated with α. The primary OA signal generated from black varnish on a petri dish exhibited 21.2dB SNR and is indicated with β, whereby reflections from the UST glass lens that arrive later in time are indicated with γ.

Fig.2g presents the averaged OA signal in the TD, when the DoV is compromised. Each LD has f_rep,j_=200KHz, N_p_=200 and a time shift between each other that results in DoV=1.875mm and t_acq_=1ms. The OA signal resulting from the simultaneous excitation using the pattern in Fig.2b is presented as a green line and is the sum of the individual OA signals obtained when each wavelength was pulsed separately (Fig.2g). We could easily detect the laser trigger interference for all wavelengths (α). The OA signals from wavelength 1 and 2 (β) are located very closely to the laser interference (α) of wavelengths 3 and 4 respectively, vastly reducing their SNR (5.7dB for wavelength 1). However, the OA signal of wavelength 3 is completely masked by the interference of wavelength 1. The reflections of the OA signal from wavelength 1, 2 and 3 (γ) are still visible. Therefore, electromagnetic interference and OA reflections further limits the DoV and SNR achieved in multiple wavelength TD optoacoustics.

Likewise, SNR limits are imposed (Fig.2h) in response to an excitation pattern (Fig.2c) that uses f_rep,j_=50kHz, N_p_=50 and DoV=7.5mm for each wavelength with t_acq_=1ms. For all wavelengths we observed that the laser interference (α), the OA signal (β) and the reflections (γ) are all visible but with lower SNR (18.8dB for wavelength 1). Finally, with f_rep_,_j_=50kHz, N_p_=200 and DoV=7.5mm for each laser the OA signal of all 4 lasers can be recorded without SNR losses (21.2dB for wavelength 1) but with t_acq_=4ms.

Conversely, FCOT operation for four wavelengths uses f_rep,1_=200kHz, δf=125Hz and N_p_=200 and recovers each signal without cross-talk between wavelengths (Fig.2j), experimentally confirming theoretical predictions. FCOT is able to provide high SNR for concurrent excitation with multiple wavelengths, without extending the acquisition time (1ms) and achieving the same DoV (7.5mm) and SNR (21.2dB for wavelength 1).

We also compared the SNR obtained in conventional FD optoacoustics to FCOT by employing the same black varnish phantom and single wavelength illumination at 445 nm (**see Suppl. FigS1**). FD optoacoustics employed a sine wave of 20 MHz frequency with adjusted mean power to equal the mean power output of the FCOT pulsed pattern used for 6.8 ns pulses at 200 KHz. FC demonstrated an SNR that was 20.8 dB higher compared to FD excitation.

### FCOT multi-wavelength imaging of tissues and tissue dynamics *in-vivo*

While the theoretical merits of FC optoacoustic operation were demonstrated with phantom measurements, a next critical step was to examine whether FC could offer realistic implementations. For this reason, we aimed to investigate whether the theoretical advantages could lead to operational characteristics (SNR, acquisition speed) that would render FCOT appropriate for in-vivo applications. A particular unknown parameter in this interrogation was the FCOT performance achieved with multiple wavelengths using laser diodes, as it would be impractical to implement FCOT with multiple solid state lasers due to cost and size. To examine the merits of using low-cost technology, we investigated the performance of multiple laser diodes to image vasculature and lymphatics *in-vivo*, using the mouse ear as a model. This imaging target was selected as it is a typical tissue where TD optoacoustic implementations have been conventionally demonstrated on in the past.

First, we assessed whether FCOT could produce high-quality images from biological specimens. We employed FC illumination at 445nm and 465nm and resolved oxy- and deoxy-hemoglobin (Fig.3a,b) based on their spectral difference, with deoxygenated hemoglobin absorbing higher at 445nm, and vice-versa, after reconstructing data collected on a grid (**see methods**). Superposition of Fig. 3a and Fig.3b revealed a color-coded composite image (Fig.3c) of the relative vascular oxygenation, with red color corresponding to higher oxygenation levels. We further confirmed that FCOT can produce depth-resolved images (**see Suppl. FigS2**) without cross-talk between the wavelengths, offering first evidence that FCOT can acquire images from biological tissues based on LDs.

**Figure 3.**
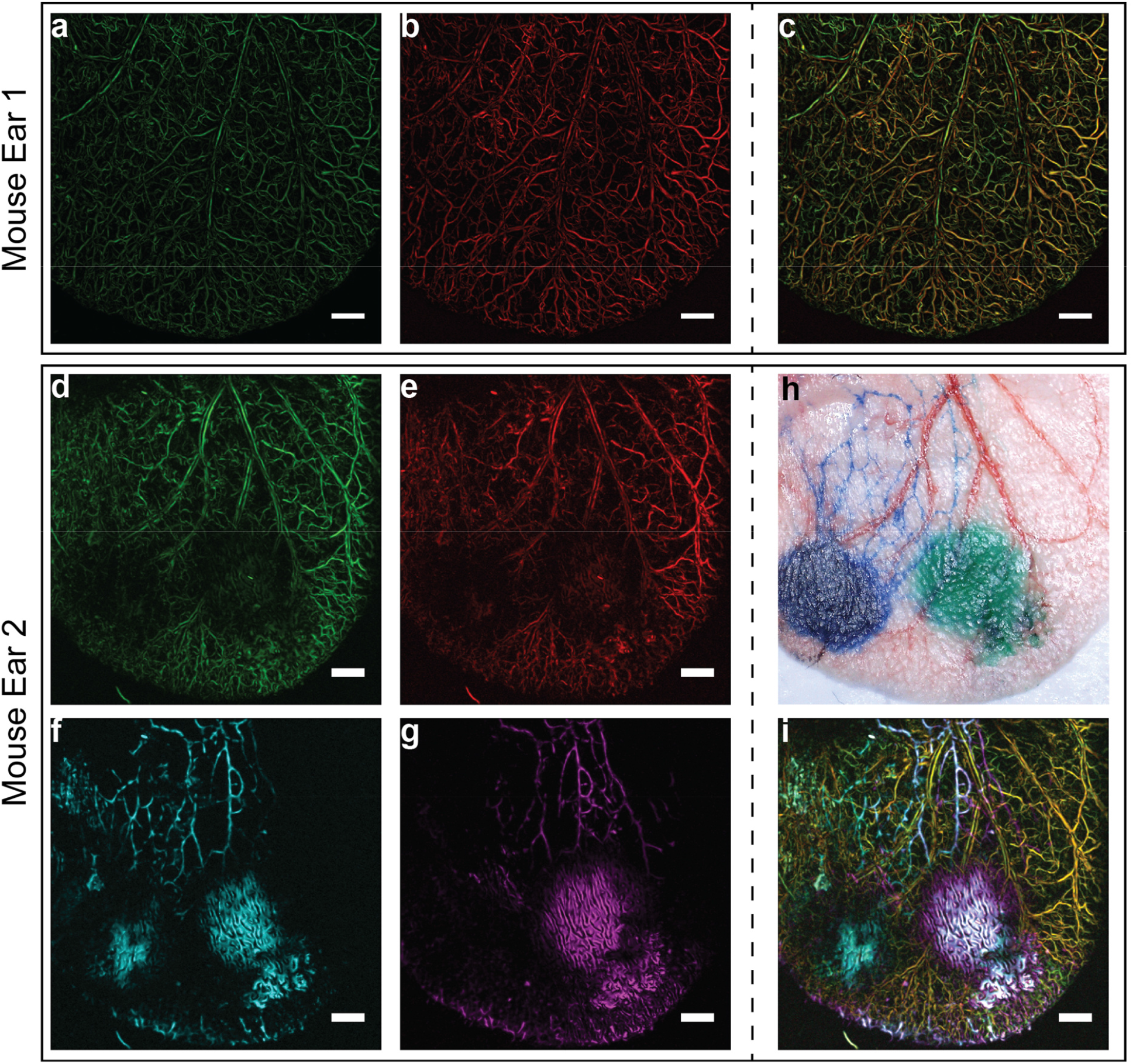
In-vivo imaging using FCOT. **a** and **b** present a mouse ear at the 2 wavelengths with high resolution. **c** shows the composite image color-coded with red indicating higher oxygenation levels compared to green. **d, e, g, h** presents a second mouse ear at the 4 wavelengths. **f** a bright field image of the mouse ear. Intradermal injection of Evan’s Blue and ICG can be seen in images **f**,**g**,**h. g** and **h**, the injected dyes enter the lymphatic vessels that present a different structure than blood vessels. **i** presents the composite image of all 4 wavelengths. We can observe oxygenated (red),de-oxygenated (green) blood vessels and lymphatic vessels after uptake of Evan’s Blue (magenta) and ICG (cyan) at the same time. All images are maximum amplitude projections of reconstructed images. Green is 445nm, Red is 465nm, Magenta is 638nm, Cyan is 808nm, scale bar 1mm.

We next aimed to identify whether FCOT could visualize multiple moieties in tissue without compromising operational characteristics as in the TD. We introduced exogenous contrast by intradermal injection of Evan’s Blue and Indocyanine Green (ICG) and applied 4-wavelength FCOT to simultaneously resolve arteries and veins (Fig.3d,e) and lymphatic vessels revealed by contrast enhancement (Fig.3f,g). Fig.3h shows the corresponding bright-field image of the mouse ear. The composite image of four wavelengths (Fig.3i) enabled visualization and co-localization of the vascular and lymphatic vessels. Notably, the use of an ultra-wideband transducer enabled to resolve the fine structures represented by the vessels as well as the large absorbing areas that were formed around the injection sites. FCOT can therefore be effectively used to perform OA imaging with multiple wavelengths simultaneously *in-vivo* with an acquisition of ∼30mins, while the same multispectral implementation in TD requires ∼2hours.

The FC acquisition acceleration over TD demonstrated in Fig.3 also points to an FCOT use for detecting rapid changes simultaneously using multiple wavelengths. We therefore applied FCOT to monitor oxygenation fluctuations in the mouse ear during an oxygen stress test *in-vivo*. Fig.4a shows the composite OA image from 2 blue wavelengths revealing the oxygenated and de-oxygenated vessels. We selected a 2mm line (Fig. 4a; blue box) to acquire signals repeatedly (continuous FC operation) at a rate of ∼4Hz. A B-scan (indicated by the letter (i) on Fig.4a) revealed an artery (red) and a vein (green) as a function of depth. The oxygen challenge was provided by alternating the composition of the breathing gas from 0.8 liters per minute (lpm) of 100% O_2_ (Fig.4b; “O_2_”) to 0.6 lpm of 20% O_2_ plus 0.2 lpm CO_2_ (Fig.4b; “Air”). Fig.4b plots the ratio of the wavelength 2 signal (S_2_) over the wavelength 1 signal (S_1_) over time. The S_2_/S_1_ ratio is indicative of the relative changes of oxygenation in the vessels over time. As expected, higher oxygenation was observed in the artery than the vein throughout the experiment. Oxygenation was reduced during the supply of air and increased when 100% O_2_ was provided. The oxygenation levels in the artery increased faster in response to the change from Air to O_2_ supply compared to the vein, revealing the expected dynamics of oxygen supply to tissues.

**Figure 4.**
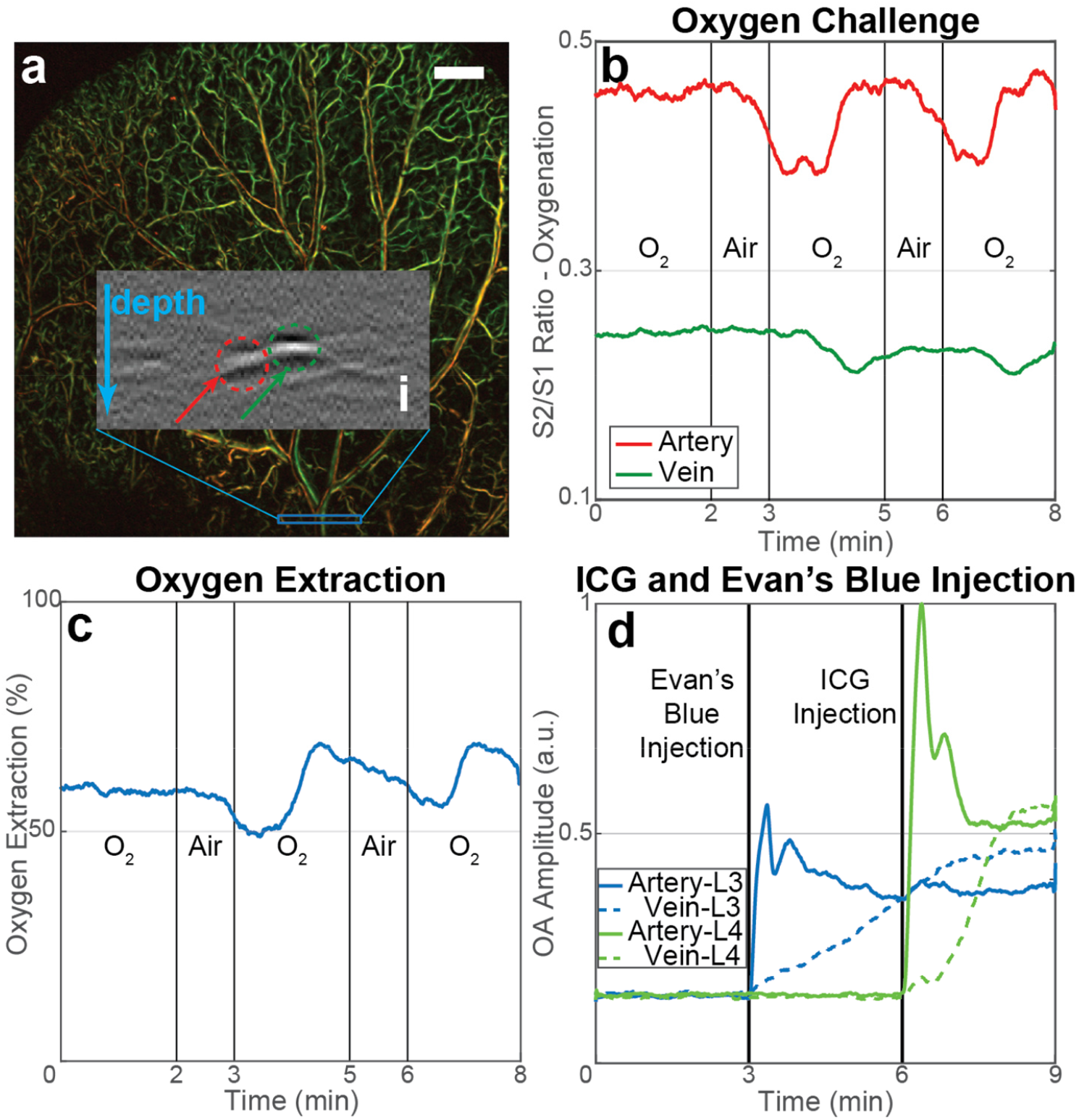
In-vivo vascular dynamics revealed by FCOT. Oxygen challenge experiment and dye injection monitoring in the central vein and artery in the mouse ear using FCOT imaging. Green is 445nm, Red is 465nm, Magenta is 638nm, Cyan is 808nm, scale bar 1mm. **(a)** shows the OA images in the blue wavelengths, showing oxygenated (red) and de-oxygenated (green) vessels. We performed continuously BScans on the blue region denoted in **(a)** and **i** shows such a cross-section. The green arrow and region in **i** indicates the selected vein, and the red arrow and region indicates the selected artery. (**b)** demonstrates the changes in the ratio between the OA signal intensity in wavelength 2 (S_2_) to that of wavelength 1 (S_1_) in time during an oxygen stress test. The oxygen saturation is proportional to the ratio S_2_/S_1_. The oxygen saturation changes faster in the artery than in the vein, as expected. **(c)** shows the oxygen extraction rate during the same experiment. **(d)** presents the signal intensity at wavelengths 3 and 4 (S_3_ and S_4_ respectively) at the same artery and vein indicated in **(a)** during intravascular injection of the 2 dyes, Evan’s Blue and ICG. In both cases the signal intensity increases first in the artery and later in the vein.

The oxygen extraction rate (OER in Fig.4c), an indication of the oxygen uptake by cells, was calculated as OER=(O_ca_-O_cv_)/O_ca_, whereby O_ca_ and O_cv_ is the oxygen saturation in the central artery and vein, respectively^27^. We observed that OER slightly dropped shortly after the Air supply period, suggesting a delayed response in cell oxygenation. When the oxygen supply was increased during the second and third O_2_ supply periods, an increased OER was observed followed by a return to normal levels, suggesting that cells consumed more oxygen as it became available. The oxygen stress-test experiment confirms FCOT as a method that can be employed in studying tissue dynamics with high localization ability, confirmed by intravenous injection of ICG and Evan’s Blue. Using the same 2mm observation field, FCOT recorded contrast agent dynamics at 638 and 808 nm (Fig.4d). We observed a similar post-injection pattern for both dyes, revealing two distinctive peaks before settling to a baseline value, indicative of the circulation dynamics of the dyes in the vascular system. We further resolved the delayed appearance of the agents in the vein for both dyes, whereby the observed signal increased at a lower rate, compared to the artery, a pattern attributed to dye diffusion in the tissue capillary network.

## Discussion

We introduce an optoacoustic method based on the frequency comb concept, which challenges the prevailing notion that TD implementations offer the best optoacoustic performance. By combining the advantages of TD excitation with FD analysis, FCOT offers superior performance to TD at multi-wavelength excitation. Increasing the number of wavelengths (N) in TD compromises the depth of view, SNR or the total acquisition time. FCOT can provide simultaneous illumination at multiple wavelengths without sacrificing any of these parameters, enabling either a higher SNR by a factor of sqrt(N) per wavelength, shorter acquisition time by a factor of N, or N times more depth of view per wavelength compared to TDOA. Moreover, FCOT provides practical implementation of an FD system, by generating multiple discrete frequencies, without the need for individual modulation and demodulation of wavelengths in each of these frequencies. In this way it avoids impractically long acquisition times or complex instrumentation.

A critical aspect herein was not only to show the theoretical superiority of the method, but also to demonstrate that it can lead to practical implementations. An enabling technology that allowed such performance was the use of overdriven continuous wave laser diodes (CW-LD)^25^ which are inherently cost-effective, portable and compact, leading to systems with the potential of high dissemination. Imaging at four wavelengths differentiated blood vasculature from lymphatic vessels *in-vivo*, reducing the acquisition time from 2 hours required in TD to ∼30 mins. FCOT also achieved a 4-fold higher acquisition rate compared to TD when performing B-scans over the main artery and vein, enabling the monitoring of relative oxygenation changes and calculation of oxygen extraction rate. Therefore, FCOT is particularly suited for dynamic measurements or for experiments that require anaesthesia, improving also the associated throughput rate over TD implementations. While the maximum number of wavelengths available in FCOT is limited by operational characteristics, up to 28 wavelengths can be simultaneously employed for f_rep,1_=200kHz, N_p_=100 and 22-78MHz, compared to only 5 wavelengths available to TD implementations with the same operating parameters and a DoV=1.5mm for each wavelength (see Supplementary material).

LDs are ideally suited for FCOT. Besides their small form factor and availability at multiple wavelengths, LDs can be pulsed at very high pulse repetition rates, matching the uniquely optimal FCOT ability to multiplex and average signals, increasing the SNR. New and more powerful LDs entering the market will contribute to further improvements of FCOT performance, offering compact and low-cost systems with high dissemination potential for various applications ^8,26^.

In summary, FCOT enables for the first time fast, high SNR imaging using multiple wavelengths simultaneously without compromising the DoV and can offer a valuable tool for studying dynamic molecular processes, revolutionizing how multispectral OA imaging will be performed in the future.

## Methods

### 4 Laser Diode Raster Scanning Optoacoustic Tomography System for FCOT

Supplementary Figure 1 presents the multispectral raster scanning optoacoustic mesoscopy (RSOM) system developed to test the advantages of FCOT. Matlab (Matlab 2016b, Mathworks, USA) was installed on a PC, controlling the system. The PC controls the dual-channel stages driver (C-867.260, Physik Instrumente, Germany) that drives the dual x-y stages placed on the Scanning Head and is synchronized with 2 Arbitrary Waveform Generators (AWG, 33522B, Keysight, USA). The AWGs trigger the laser diode drivers and provide a synchronization pulse to the Data Acquisition Card (DAQ, 12-bit, 200MS/s, Razor Express 14×2 Compuscope, Dynamic Signals LLC, USA) for synchronous acquisition of the signal. The light output of the Illumination system is directed into the Scanning Head, with a small portion entering a photodiode (DET10A2/M, Thorlabs, USA). The signal from the photodiode is analog filtered (BLP-90+, Minicircuits, USA) and recorded to monitor the pulse to pulse energy fluctuations and time jitter, corrected for both during signal post-processing. The OA signal from the UST is amplified with a 60dB gain amplifier (Miteq AU-1291-R, Miteq, USA) and analog filtered (BLP-90+ and ZFHP-1R2-S+, Minicircuits, USA) before being digitized by the DAQ to avoid aliasing.

The Illumination system consists of 4 CW laser diodes that were overdriven with 4 high-current, short-pulse laser diode drivers developed previously^25^. The laser diodes used in this work are the LDM-445-6000 (LaserTack, Germany) emitting at 445nm, the LDM-465-3500 (LaserTack, Germany) emitting at 465nm, the HL63283HG (Ushio, Japan) emitting at 638nm and the K808D02FN (BWT, China) emitting at 808nm, named laser 1, laser 2, laser 3 and laser 4 respectively. Each laser diode is focused in a multimode fiber. In order to position each laser diode in a manual X-Y stage (CXY1, Thorlabs, USA), a collimating lens (C340TMD, Thorlabs, USA) is placed in front of it on a manual z-stage (SM1Z, Thorlabs, USA), followed by a focusing lens (C560TME, Thorlabs, USA) that is kept stable and the fiber on a x-y stage (CXY1, Thorlabs, USA). The fiber with a 200um core diameter and 0.22NA was one of the 4 inputs of 4×4 fiber power combiner. The 4 outputs of the fiber combiner (MPC-4-M21-M41-P23, Lasfiberio, China) contain ∼25% of the input power of each input fiber and are also multimode fibers with a 200um fiber core and 0.22NA. One of the outputs is connected to a custom made 95-5% splitter (LTL 500-93310-95-1, LaserComponents Germany GmBH, Germany) and the 5% fiber was connected to the photodiode. The 3 outputs of the power combiner and the 95% fiber of the splitter were terminated with 1.25mm ferrules (SFLC230, Thorlabs, USA) and directed to the Scanning Head.

The Scanning Head consists of the x-y stage (U-723 XY, Physik Instrumente, Germany), the 3D printed holder, the ultrasound transducer (UST, HFM23, Sonaxis, France) with a 50MHz central frequency and 112% relative bandwidth, 3mm focal length and 0.5NA, and the 4 output fibers arranged in a circular pattern around the UST. The output of the 4 fibers is designed to cross at the focal spot of the UST to achieve a maximum energy density on the sample.

Scanning and recording is done in a sweeping-like motion using the stages driver as the master in the system. The x-stage moves in a straight line with constant velocity. When the x-stage has traveled a specific distance, equal to the step size, the stages driver sends a signal to the AWGs to trigger the laser diodes. Each laser diode is triggered using a pulse train with repetition rate f_rep1_-f_rep4_ for each laser diode and N_p1_-N_p4_ number of pulses at each point (A-Scan) in the B-Scan. After the stage has traveled the desired distance, the B-Scan is completed and the stages stop moving. The y-stage is moved to the next y-position and the x-stage can now perform the next B-Scan in the opposite direction. For the FCOT excitation we used f_rep,1_=200000Hz, f_rep,2_=200125Hz, f_rep,3_=200250Hz and f_rep,4_=200375Hz and N_p1_=100 and N_p2_=N_p3_=N_p4_=101 pulses for lasers 1, 2, 3 and 4 respectively. For a detailed derivation of these values see Supplementary Material.

To compare the FD OA to FCOT excitation we used a fiber-coupled 450-nm laser diode (FBLD-450-0.8W-FC105-BTF; Frankfurt Laser Company, Germany) connected to an analog laser driver (BFS-VRM 03 HP; Picolas, Germany). The output fiber of the laser was pumped into one of the input fibers of the 4×4 fiber combiner, so that the illumination of the sample is identical for the FD excitation and the FCOT excitation system.

### Reconstruction algorithm

Image acquisition occurs in a large Field of View (10×10 mm^2^) and with a scanning step size of 10um in order to be much lower than the lateral resolution of the system, which is calculated to be 38um. Time-resolved signals detected at each scanning position correspond to the integration of acoustic spherical waves originating from the illuminated optical absorbers within the detection angle of the transducer. Therefore, unprocessed images obtained directly from the system are heavily blurred, and require further processing to obtain high-contrast and high-resolution images. To do so, a back-projection algorithm is implemented in the Fourier domain^28^ to recover an (acoustically) diffraction limited image. The resulting image is then corrected by the system impulse response^29^ and further processed with a vesselness filter for display^30^. The raw data of a single B-Scan do not allow for 3D image reconstruction. Therefore, a simplified version of the back-projection algorithm was developed, which operates in two dimensions and approximates the transducers sensitivity field as a conic section along the B-Scan direction only.

### Laser Diode emission spectra, hemoglobin and dye absorption spectra

We recorded the emission spectra of the 4 laser diodes used in the multispectral laser diode RSOM with a spectrometer (USB4000, OceanOptics, UK) and the peak emission wavelengths were 444.3, 460.1, 636.8, 804.9 nm with a variance of 1.6, 1.7, 1.9, 2.2 nm respectively and an R^2^ confidence level higher than 0.96 for all cases.

Hemoglobin has a broad absorption spectrum over the visible and near infrared range with a higher absorption at the lower wavelengths of the spectrum. The absorption of de-oxygenated hemoglobin is higher than the absorption of oxygenated hemoglobin at 444nm. The absorption of oxygenated hemoglobin is higher than the absorption of de-oxygenated hemoglobin at 460nm. The total absorption of hemoglobin at wavelengths 1 and 2 is much higher than that at wavelengths 3 and 4. Due to the low energy output of the laser diodes, this is enough to assume that we only detected OA signals from hemoglobin at wavelengths 1 and 2. This has been confirmed from the *in-vivo* mouse ear experiments. Moreover, we can estimate the relative changes of oxygen saturation^31^ by calculating the ratio of the signal intensity of wavelength 2 over that of wavelength 1, S_2_/S_1_, with a higher ratio indicating a higher oxygen saturation.

To induce contrast and increase the SNR at wavelengths 3 and 4 we used two dyes, Evan’s Blue (Sigma-Aldrich, Germany) and ICG (VERDYE, Germany). Evan’s Blue has a peak absorption at 640nm and a minimum at 740nm^32^ making it the appropriate dye to enhance the contrast for wavelength 3, since this is the only wavelength where we can detect OA signals. ICG in blood plasma has a peak absorption at around 810nm and a low absorption at 637nm^33^, making it appropriate to induce OA contrast at wavelength 4 with minimal contrast at wavelength 3 and no contrast at the other wavelengths, confirmed by the mouse ear experiments.

### Maximum Permissible Exposure (MPE) Limits Compliance

The energy per pulse on samples was measured with a stabilized thermal power meter (PM160T, Thorlabs, USA) and calculated as 189, 137, 142, 153 nJ per pulse for lasers 1, 2, 3, 4 respectively. The pulse width was estimated as 6.7, 6.7, 10.2, 10.2 ns full-width-at-half-maximum (FWHM) for each laser respectively. Using a USB CCD camera (daA1920-30um; Basler AG, Germany) the illumination spot on the surface of the sample was measured to be a circle with a diameter of ∼1mm. Using the above mentioned repetition rates for each laser diode and a scanning speed of 10mm/s for a scanning step size of 10um we can calculate the sample exposure.

The total exposure of the sample for simultaneous illumination with all four wavelengths is calculated to be 19.8uJ/cm^2^ per pulse and 3.96W/cm^2^ mean exposure, well below the MPE limits of 20mJ/cm^2^ and 18W/cm^2^, imposed by the American National Standards Institute^34^.

### Mouse handling and Imaging Protocol

For our experiments we employed two 5 to 6-week old female Athymic nude-Foxn1^nu^ mice (Envigo, Germany). During all measurements, the mice were anesthetized by 1.6% Isoflurane (cp-Pharma, Germany) with 0.8lpm carrier gas flow and kept on constant body temperature with a red lamp and a heating plate. All procedures involving animal experiments were approved by the Government of Upper Bavaria, under animal protocol number 55.2-2532.Vet_02-18-120.

The first mouse was used for the experiments presented in Fig 3. The lymphatic ear vessels were highlighted by intradermal administration of 5ul ICG (5mg/ml) and 5ul Evan’s blue (1%)into the ear tip of the second mouse. ICG was administered ∼30 min and Evan`s blue ∼10 min before imaging start to ensure the lymphatic drainage of the dyes.

A second mouse was used for the experiments demonstrated in Fig 4. For the oxygen stress experiment, we supplied different isoflurane carrier gas combinations or breathing conditions through a nose mask. The mouse was breathing alternatively 0.8 lpm of 100% oxygen (O_2_) and a combination of 0.6 lpm medical Air (20% Oxygen) plus 0.2 lpm carbon dioxide CO_2_ (Air). For the dye diffusion experiment we initially acquired the background data and after 3 minutes intravenously injected 100ul of 1% Evan`s Blue solution and after another 3 minutes 100ul of the 5 mg/ml ICG solution. Both mice were sacrificed immediately after imaging.

## Data Availability Statement

The authors declare that the data supporting the findings of this study are available within the paper and its supplementary information files.

## Code Availability Statement

The authors declare that for data collection the commercially available software from GaGe (Dynamic Signals LLC, USA) and Matlab 2016b (Matlab, Mathworks, USA) was used. Data analysis was conducted in Matlab using its built-in functions. The software code is available from the corresponding author upon reasonable request.

## Acknowledgements

The research leading to these results has received funding by the Bundesministerium für Bildung und Forschung (BMBF), Bonn, Germany (Project Sense4Life, 13N13855), from the European Research Council (ERC) under the European Union’s Horizon 2020 research and innovation programme under grant agreement No 694968 (PREMSOT) and from the European Union’s Horizon 2020 research and innovation programme under grant agreement No 732720 (ESOTRAC). We would like to thank Dr. Sergey Sulima for his help writing the manuscript.

## Supplementary Material

**Supplementary Figure1.**
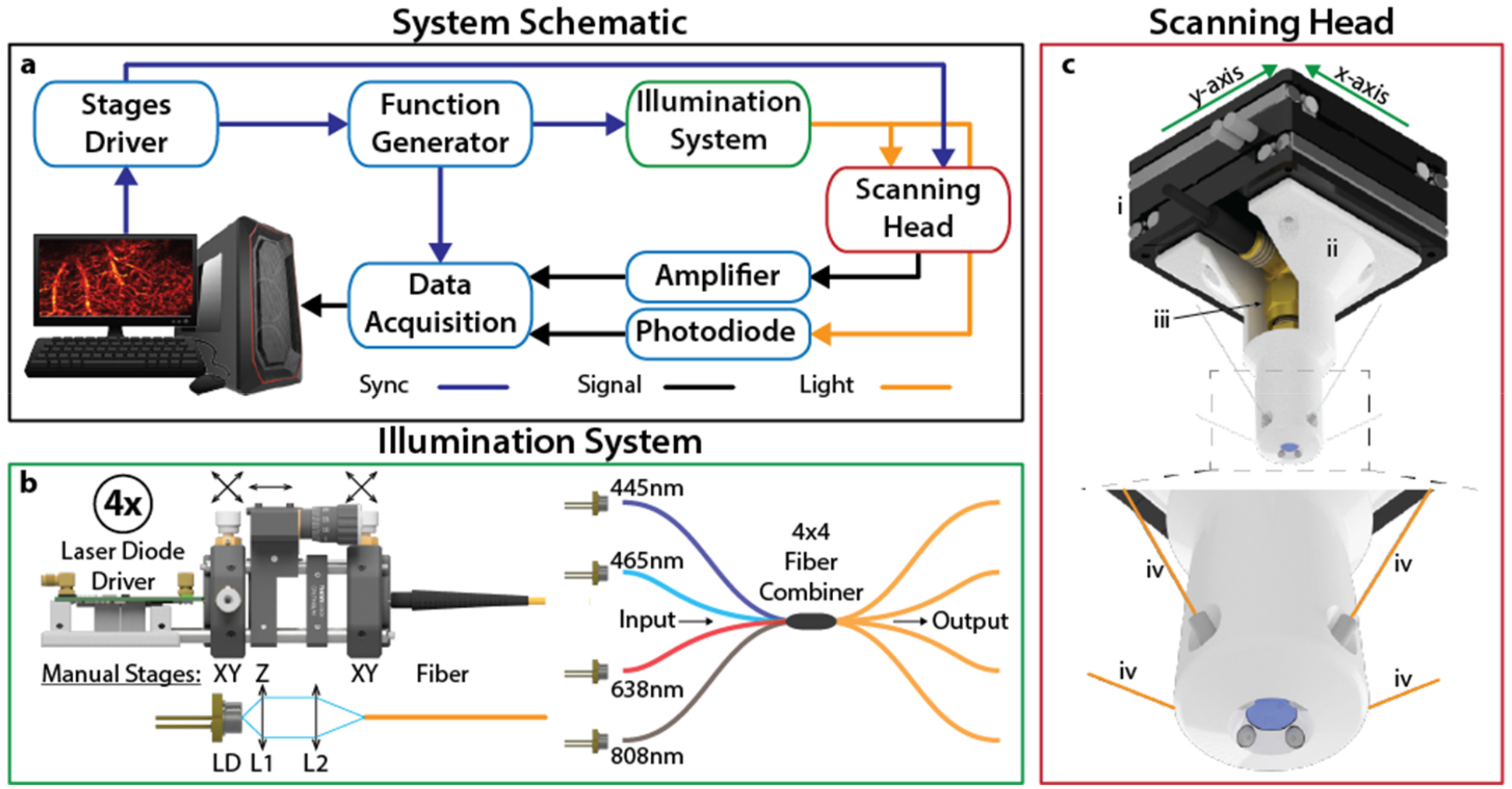
Multispectral Raster Scanning Optoacoustic Mesoscopy system using 4 laser diodes. **a** The schematic of the system developed showing the electrical and optical connections of the different parts of the system. **b** The laser diode illumination system. Each laser diode is attached to a separate laser diode driver and focused into a multimode fiber with a 2 lens system. The 4 laser diodes are coupled in a 4×4 fiber power combiner and each output of the combiner has ∼25% of the power of each input, combining all the wavelengths. **c** The scanning head of the RSOM system consisting of the x-y scanning stages (i), the 3D printed holder (ii), the UST (iii) and the 4 output fibers of the fiber power combiner (iv) arranged in a circular pattern around the UST.

**Supplementary Figure2.**
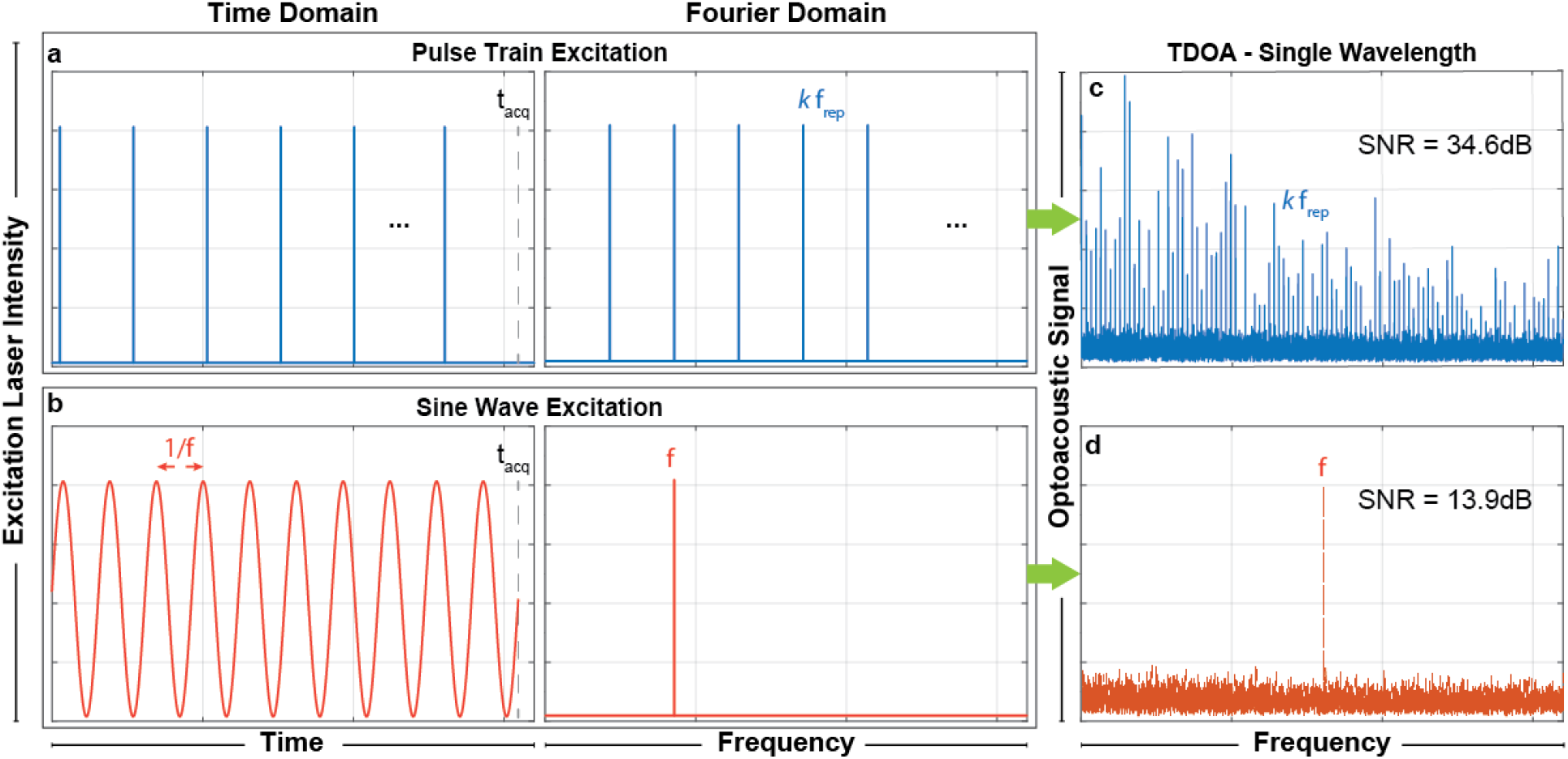
FCOT advantages over FD-OA at single-wavelength illumination. **a** The FC pulse train excitation in Time Domain and Fourier space. **b** The sinusoidal FD excitation at frequency f in Time Domain and Fourier space. **c** The optoacoustic signal from a black varnish layer on a petri-dish after excitation with the pattern in **a** in the Fourier space after recording it in time and performing the Fourier Transform. The signal appears as many discrete frequency peaks that are all harmonics of the laser pulse repetition rate f_rep_. The total OA signal SNR is calculated to be 34.6dB. **d** The optoacoustic signal from the sinusoidal excitation in **b** in Fourier space from the same sample. Now the signal appears in only 1 frequency, f, with an SNR of 13.9dB.

**Supplementary Figure 3.**
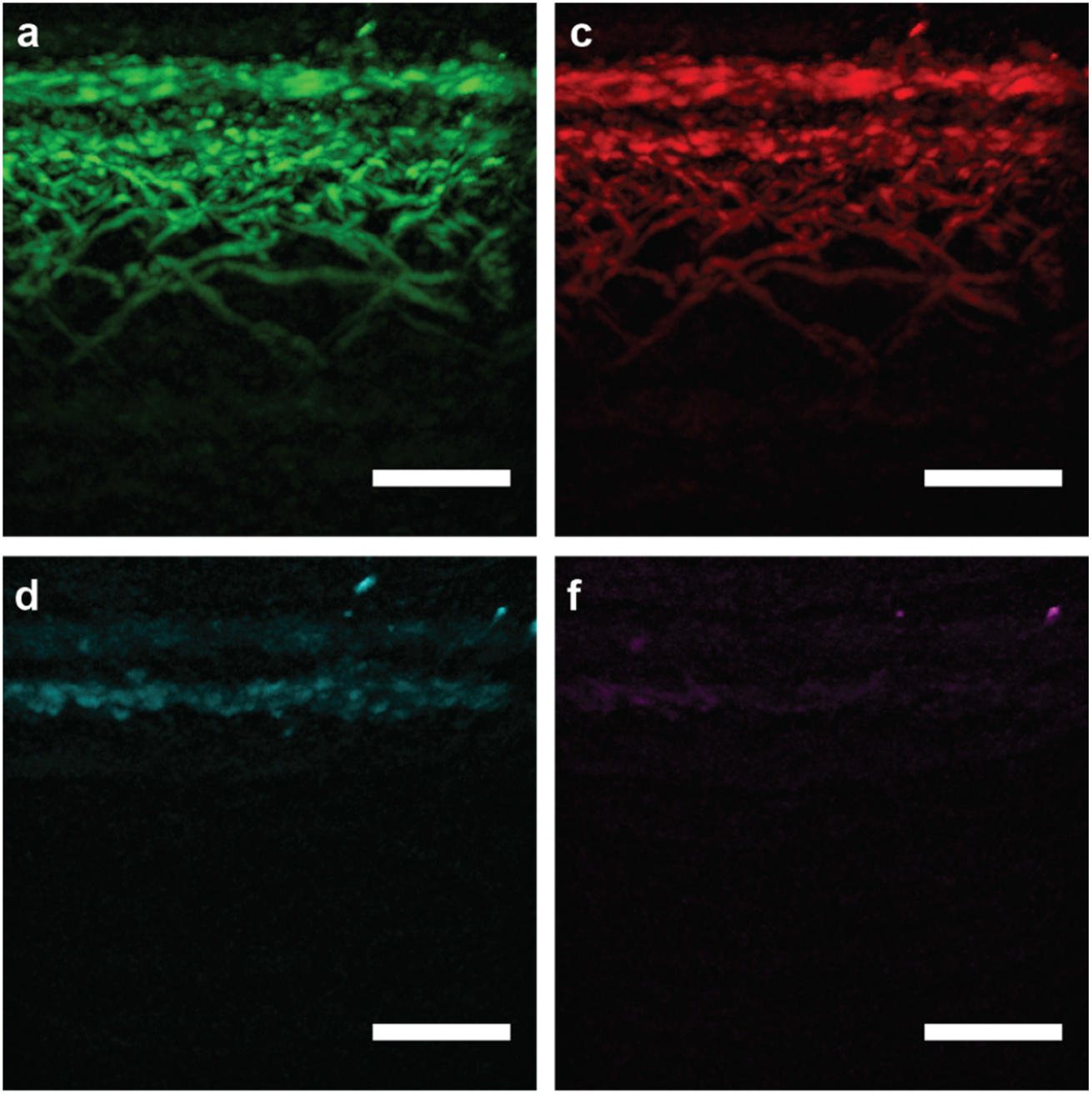
Cross-sectional in-vivo images of human skin using FCOT at four wavelengths. **a** presents the human skin at 445nm and **b** at 465nm. Both wavelengths were able to detect melanin and blood vessels located deeper in the skin. **c** presents the skin at 638nm where only weak contrast from melanin is detected. **d** shows the skin as detected at 808nm where blood and melanin absorption are weak to generate a detectable OA signal. Green is 445nm, Red is 465nm, Magenta is 638nm, Cyan is 808nm, scale bar 1mm in the horizontal direction, vertical range is 1mm total.

### Human Skin Experiment

One of the authors volunteered for this study to have their hand imaged. After consultation with the TUM Ethics Commission, no formal ethics approval was necessary. Informed consent from the participant was obtained and archived.

### Frequency Comb Algorithm – Formula derivation

In FCOT the recorded signal over N_p_ pulses is *S*_*r*_(*t)* = *S*(*t)* + *n*(*t)*, where *S*(*t)* is the periodic OA signal and *n*(*t*) the additive white Gaussian noise. Since *S*(*t*) is periodic, we can express its k-th period as *S*_*r,k*_(*t)* = *S*_*r*_(*t)* * *δ*(*t* + *kT)*, where * denotes the convolution with the delta function. From Fourier theory we know that

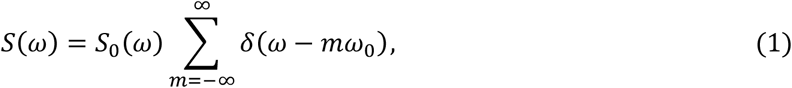

with *ω*_0_ = 2 *π*/*T* and *T* the period of *S*_*r*_(*t*). This equation indicates that the Fourier transform of a periodic signal is the Fourier transform over one period of the periodic signal multiplied by a series of deltas that confine the Fourier transform of the periodic signal to the harmonics of its repetition rate. We can now calculate the averaged signal

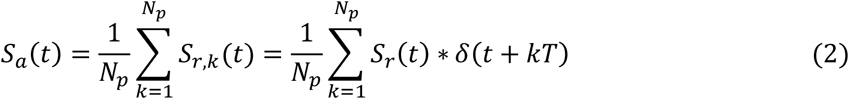

by shifting the recorded signal back by a multiple of the period each time and summing, with *t* being 0 < *t* < *T*. We can now apply the Fourier transform in equation (2):

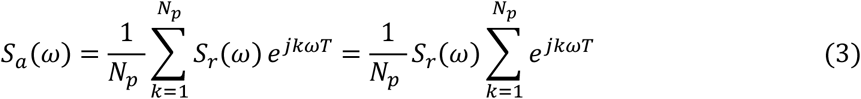

Since we know that *S*_*r*_(*t*) contains a periodic signal with additive noise we can restrict it to the harmonics of the periodic signal, i.e. multiply equation (3) by 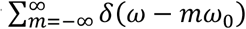:

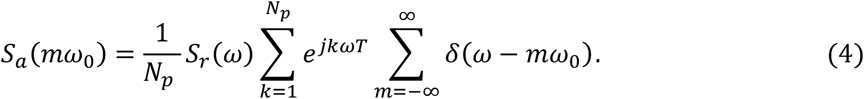

However, 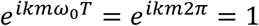 for every integer *k* and *m*. Therefore, the sum of the exponentials is *N*_*p*_ and equation (4) becomes

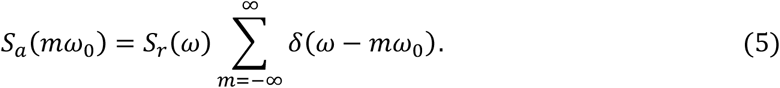

This equation tells us that the Fourier transform of the averaged signal is the Fourier transform of the recorded signal confined to the harmonics of its repetition rate. We can further expand the recorded signal to obtain

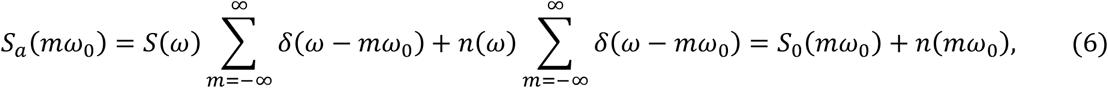

which tells us that the Fourier transform of the averaged signal is the Fourier transform of the first period of the periodic signal, what we wanted to recover, plus the Fourier transform of the white Gaussian noise confined to the harmonics of the repetition rate of the periodic signal. Since it is not periodic, the noise is significantly suppressed with its power reduced by sqrt(Np).

### Modulation parameters for the FCOT – Formula Derivation

In this work, the FCOT signal is acquired in the Time Domain using a high speed digitizer. In FCOT the N wavelengths have a different repetition rate, with a small frequency shift δf<<f_rep_. For each wavelength, we follow the same FCOT signal processing algorithm for a single wavelength, choose only the harmonics of the laser repetition rate and apply the inverse Fourier Transform to recover the TD signal. For the case of multiple wavelengths all harmonics from any laser repetition rate should be accurately resolved from all harmonics of each other’s laser repetition rate. If any of the harmonics are not well-resolved, there will be a cross-talk between the signals from different wavelengths. Therefore, the choice of the number of pulses in the pulse train and the frequency shift δf is critical to be able to separate the signal from each wavelength. In the following we present the analytical derivation of the formulas that impose the limits on the number of pulses in the pulse train for each wavelength and the frequency shift.

The first wavelength has a reference repetition rate f_rep1_, the second wavelength a repetition rate f_rep2_=f_rep1_+δf and the N^th^ wavelength a repetition rate f_repN_=f_rep1_+(N-1)*δf. The pulse train of the first laser contains N_pulses_ pulses; this leads to t_acq_=N_pulses_/f_rep1_ acquisition time. For each AScan we apply a Fourier Transform on the data to get the discrete frequency peaks in the Fourier Domain. In Fourier Domain the maximum frequency resolution will be df=1/t_acq_. The Ultrasound Transducer (UST) exhibits a specific detection frequency range from f_low_ to f_high_, taken as the -6dB cut-off point of its frequency response.

For the reference wavelength, laser 1, we need to find the harmonics that are located and the limits of the UST detection range, i.e. the first k_s_ for which, k_s_*f_rep1_>f_low_, and the last k_f_ for which, k_f_*f_rep1_<f_high_. This happens for k_s_=ceil(f_low_/f_rep1_) and k_f_=floor(f_high_/f_rep1_), where floor(x) rounds x to the nearest integer smaller than or equal to x and ceil(x) rounds x to the nearest integer greater or equal than x.

At the lower end of the UST detection range, the harmonic k_s_ of laser 1 will be very close to the k_s_ harmonic of laser 2, i.e. Δf = k_s_*f_rep2_ - k_s_*f_rep1_ = k_s_*(f_rep1_ + δf) - k_s_*f_rep1_ = ks*δf. To be able to resolve these two frequencies Δf>df, i.e. δf>df/k_s_. This condition applies a minimum limit to the frequency shift δf, with δf_min_=df/k_s_.

At the higher end of the UST detection range, the k_f_ harmonic of laser 1 will be very close to the k_f_-1 harmonic of laser N, i.e. Δf = k_f_*f_rep1_ – (k_f_-1)*f_repN_ = k_f_*f_rep1_ – (k_f_-1)*[f_rep1_ + (N-1)*δf] = f_rep1_ – (k_f_-1)*(N-1)*δf. To be able to resolve these two frequencies Δf>df, i.e. δf<(f_rep1_-df)/[(k_f_-1)*(N-1)]. This condition applies a maximum limit to the frequency shift δf, with δf_max_=(f_rep1_-df)/[(k_f_-1)*(N-1)].

We observe that as N_pulses_ decreases, the acquisition time decreases and the frequency resolution df increases. Therefore, as N_pulses_ decreases, δf_min_ increases and δf_max_ decreases. As δf_min_ should be smaller than δf_max_, this imposes a minimum limit for N_pulses_ in the case that δf_min_=δf_max_, N_pulses_min_=ceil{[(k_f_-1)*(N-1)+ks]/k_s_}. Any number of pulses in the pulse train larger than N_pulses_min_ is valid and allows for a valid definition of a range for the small frequency shift δf, between δf_min_ and δf_max_.

The minimum number of pulses depends on the reference repetition rate, f_rep1_, the UST detection range, f_low_ and f_high_, and the number of wavelengths used, N. Once a valid N_pulses_ is chosen, the range of valid δf’s then depends on f_rep1_, f_low_, f_high_, N and N_pulses_ and a valid frequency shift can be chosen.

Following the above, for the bandwidth of the UST that we used (22-78 MHz) we use a reference repetition rate of f_rep1_=200KHz and 4 wavelengths. This results in a minimum possible N_pulses_ equal to 12. Since we are using laser diodes that usually provide low SNR, we chose N_pulses_=100 to increase the SNR through averaging. For this number of pulses, δf can be in the range between 18.2Hz and 169.7 Hz. For the frequency shift we chose δf=125Hz, which gives the base repetition rate for each laser as f_rep1_=200000Hz, f_rep2_=200125Hz, f_rep3_=200250Hz, f_rep4_=200375Hz. Moreover, the number of pulses in each pulse train is adjusted to fit as many pulses as possible in the acquisition time defined by the reference repetition rate and N_pusles1_. In this case, the following N_pulses_ are chosen for each wavelength respectively, N_pulses1_=100, N_pulses2_=101, N_pulses3_=101, N_pulses4_=101.

### Wavelength limit in FCOT compared to TD OA

There is a maximum number of wavelengths that can be pulsed simultaneously in FCOT. In all cases, δf_min_ should be smaller than δf_max_, i.e. δf_min_<δf_max_, and by substituting their values as calculated above and solving for the maximum number of wavelength N_max_ we get N_max_=floor{[(f_rep,1_-df)*k_s_]/[(k_f_-1)*df] +1}. Assuming an f_rep,1_=200kHz, N_p_=100 and a -6dB UST bandwidth between 22 and 78 MHz (f_low_=22MHz and f_high_=78MHz) we get N_max_=28. For the same parameters in TD OA a repetition rate of 200kHz results in a total DoV of 7.5mm and assuming that we need at least 1.5mm DoV for each wavelength no more than 7.5/1.5=5 wavelengths can be multiplexed without compromising SNR or the total acquisition time.

